# Seasonal influenza circulation patterns and projections for 2016-2017

**DOI:** 10.1101/076638

**Authors:** Trevor Bedford, Richard A. Neher

## Abstract

This is not meant as a comprehensive report, but is instead intended as particular observations that may be of relevance. Please also note that observed patterns reflect the GISAID database and may not be entirely representative of underlying dynamics. All analyses are based on the nextflu pipeline [1] with continual updates posted to Correspondence:http://nextflu.org.

## A/H3N2

Despite the late-season 3c3.a epidemic in the USA, we predict clade 3c2.a viruses will continue to predominate in the H3N2 population. Within clade 3c2.a, the 171K variant has spread rapidly so that the majority of recent H3N2 infections are comprised of 171K viruses. Barring the emergence of a new antigenic variant, we believe 171K will continue to predominate.

We base our primary analysis on a set of viruses collected between Sep 2014 and Aug 2016, comprising approximately 100 viruses per month where available and seeking to equilibrate sample counts geographically where possible (Fig. 1). This equilibration attempts to collect equal samples from Africa, China, Europe, Japan/South Korea, North America, Oceania, South America, South Asia, Southeast Asia and West Asia. In the following analysis we collapse samples from China, South Asia, Southeast Asia, Japan and Korea into a single region referred to here as “Asia”, resulting in Asia possessing greater sample counts than North America or Europe. The only month that significantly departs from equitable sampling is Aug 2016 with 38 viruses, primarily from Europe and North America. We subsample to 100 viruses per month and not more to keep sample counts as equitable as possible across space and time. Repeating mutation and clade frequency calculations with up to 1200 viruses per month yields similar results (see below).

**Figure 1.**
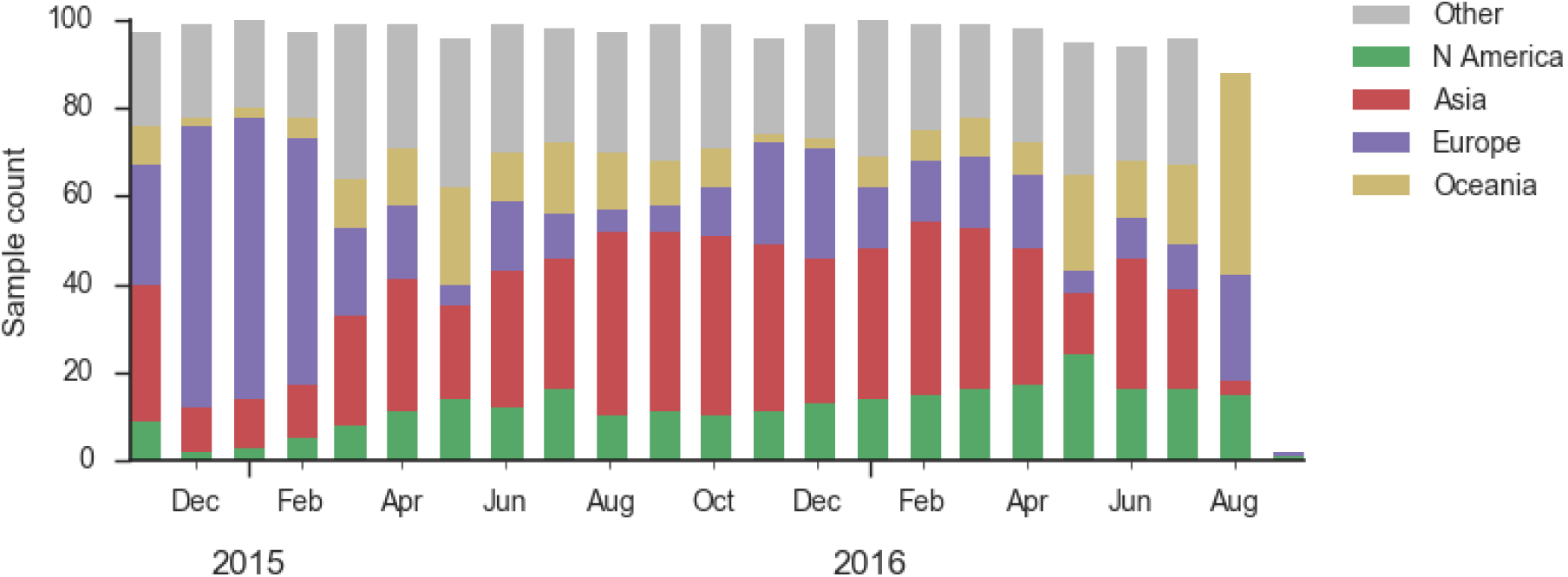
Sample counts through time and across regions. This is a stacked bar plot, so that most months there are *∼*100 total samples and *∼*15 samples each from North America and from Europe.

Viral clades 3c3.a, 3c3.b and 3c2.a emerged from the Texas/2012 background in early 2014 and rapidly spread through the viral population. Subsequently, we have observed competition among these clades, with 3c2.a viruses being globally dominant beginning in 2015 (Fig. 2). Recently, we have observed the steady decline of 3c3.b viruses. At this point, we estimate that they are either extinct or nearly extinct. 3c3.a viruses were largely replaced by 3c2.a viruses starting in 2015. However, we have observed an anomalous late-season epidemic of 3c3.a viruses within the USA from Jan to Aug 2016. Elsewhere in the world, 3c2.a viruses have remained dominant, particularly in Asia, where we estimate 3c2.a frequencies to be *>*95% throughout 2016.

**Figure 2.**
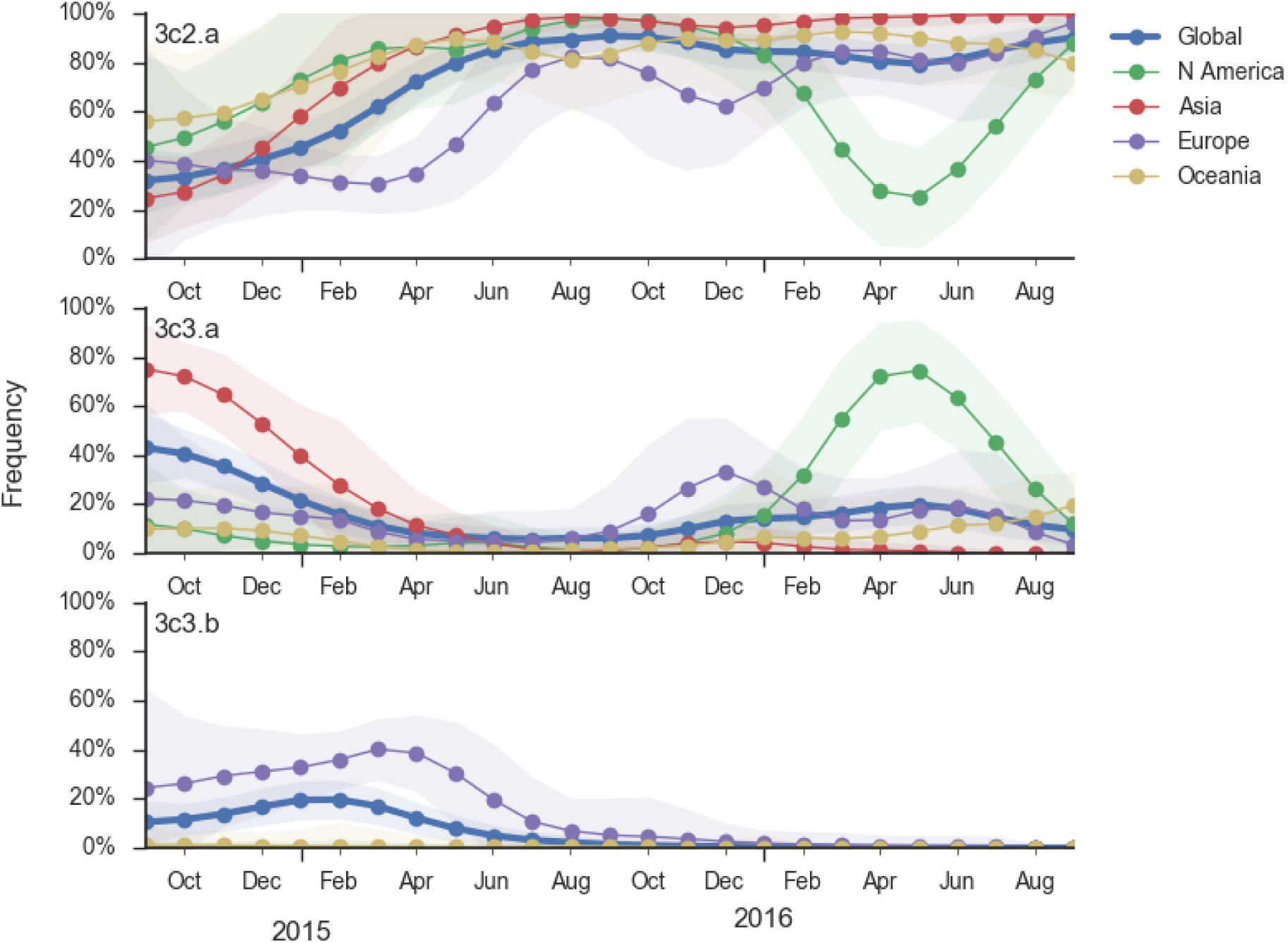
Frequency trajectories of H3N2 clades. We estimate frequencies of different clades based on sample counts and collection dates. We use a Brownian motion process prior to smooth frequencies from month-to-month. Transparent bands show an estimate the 95% confidence interval based on sample counts. The final point represents our frequency estimate for Sep 1 2016.

The late-season 3c3.a epidemic in the USA is puzzling. However, we suspect that this epidemic is due to epidemiologic circumstance rather than the emergence of a selective variant. The strongest evidence for this is that there is not a single clade within 3c3.a that is spreading throughout the USA (Fig. 3). Instead, a variety of 3c3.a viruses are spreading, each with different HA1 mutations. The emergence and spread of an adaptive variant would have appeared as a single clade bearing a characteristic epitope mutation. This is not what we see. One small clade, however, carries the S145N and F193S mutations close to the receptor binding site which resulted in antigenic evolution in previous years. This is a small clade restricted to viruses from US and remains at less than 1% global frequency.

**Figure 3.**
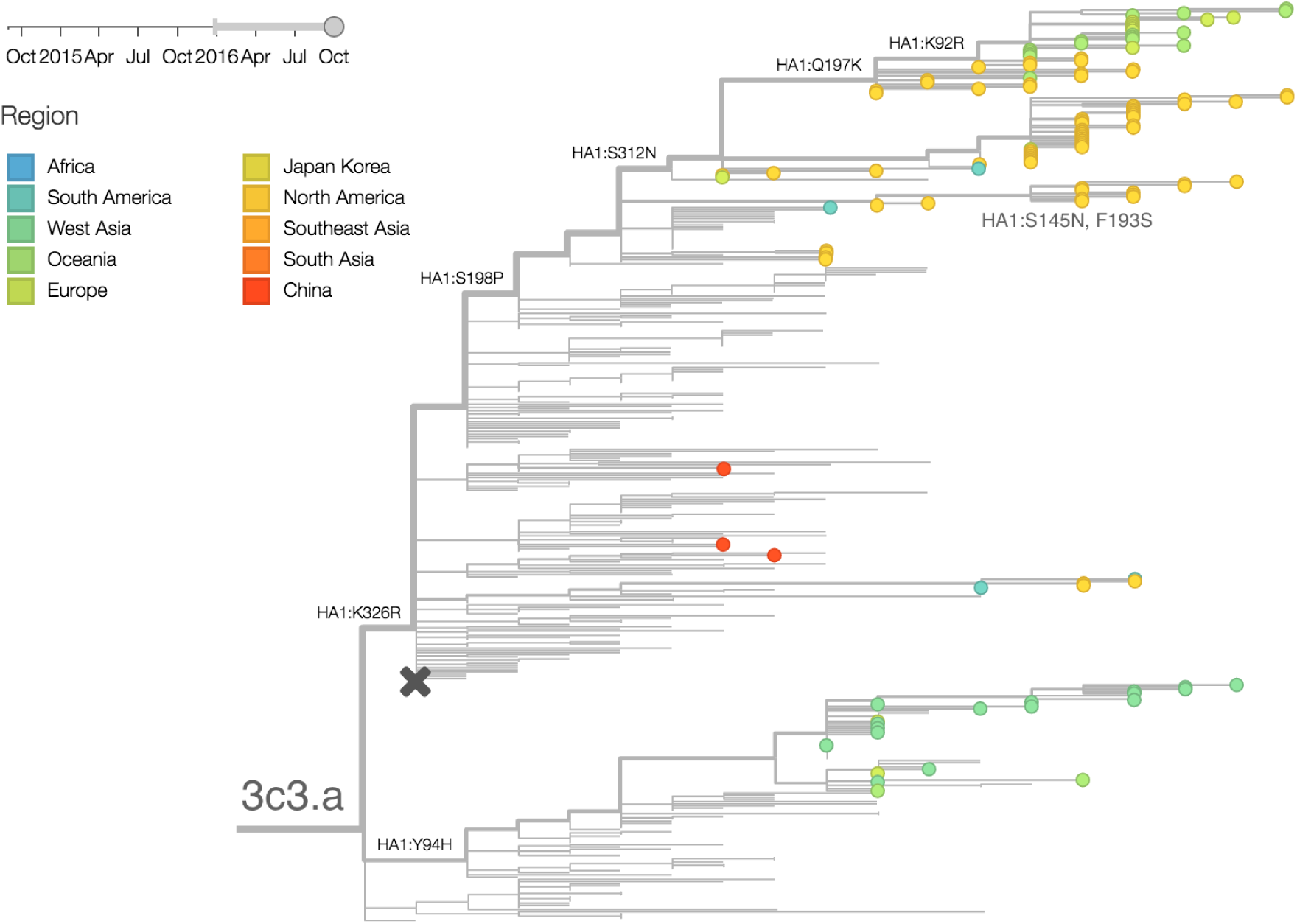
H3N2 / 3c3. a phylogeny colored by geography.

We doubt that these 3c3.a viruses will spread globally. It is conceivable however, that the 2016-2017 USA season could derive from over-summering transmission chains. This would be an unusual event, but not unheard of [2,3]. In one recent season (2008-2009), USA viruses derived primarily from the previous USA season. In other years (at least back to 2000), USA seasons were primarily reseeded from elsewhere (Fig. 4). We believe that the 2016-2017 USA season will likely derive from 171K viruses due to their rapid spread (see below).

**Figure 4.**
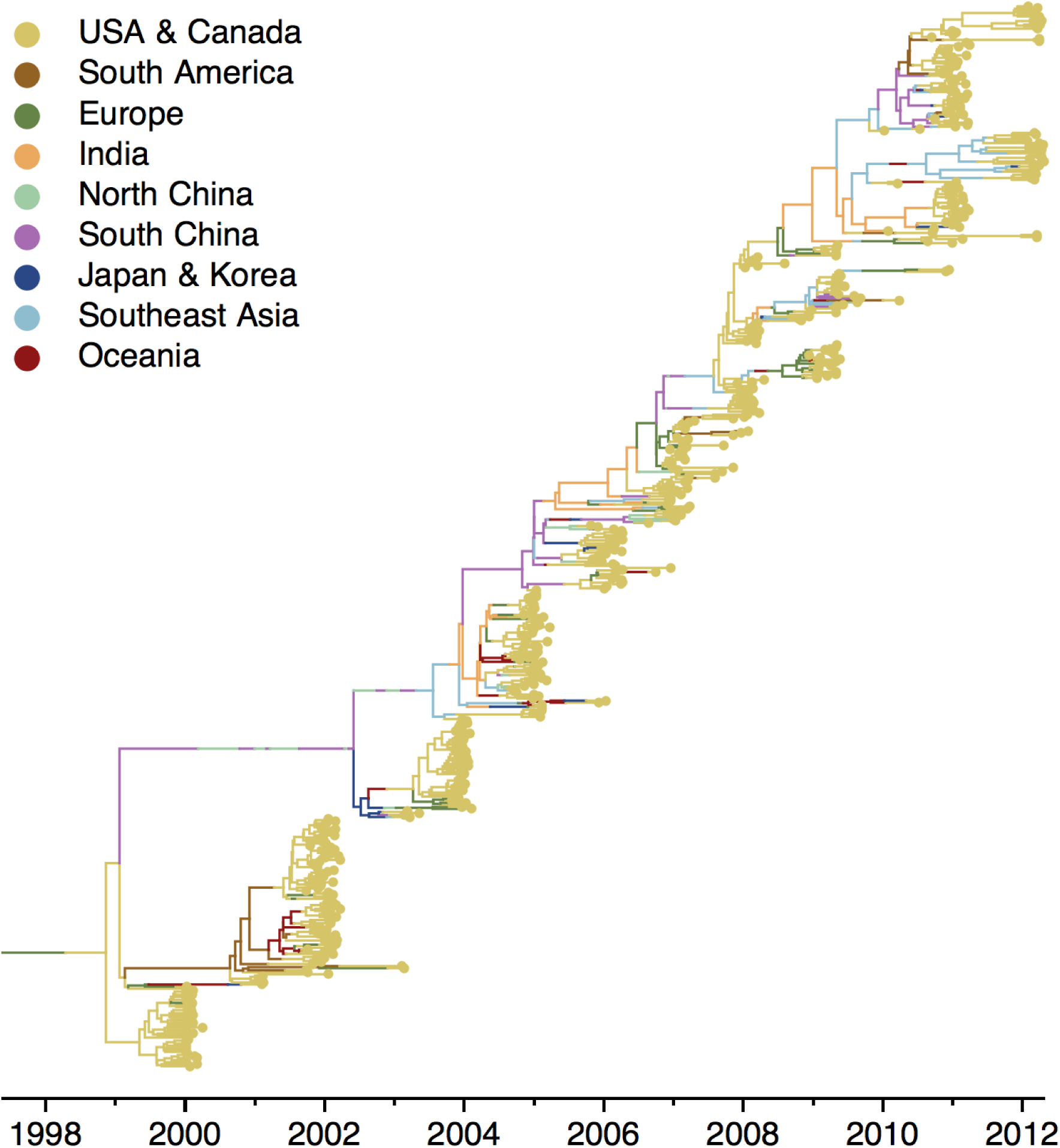
Historical phylogeny of H3N2 viruses from North America from Bedford et al. (2015) [3].

The dominant 3c2.a viruses have continued to genetically diversify with the emergence of multiple subclades (Fig. 5). We observe 3 subclades of decent frequency in 2016 viruses. These are characterized by HA1 mutations 142K/197R, HA1 mutation 197K and HA1 mutation 171K + HA2 mutations 77V/155E. The 171K variant comprises approximately 58% of 3c2.a viruses collected in 2016. Within 171K, the variant 121K has emerged and comprises approximately 23% of 3c2.a viruses collected in 2016.

**Figure 5.**
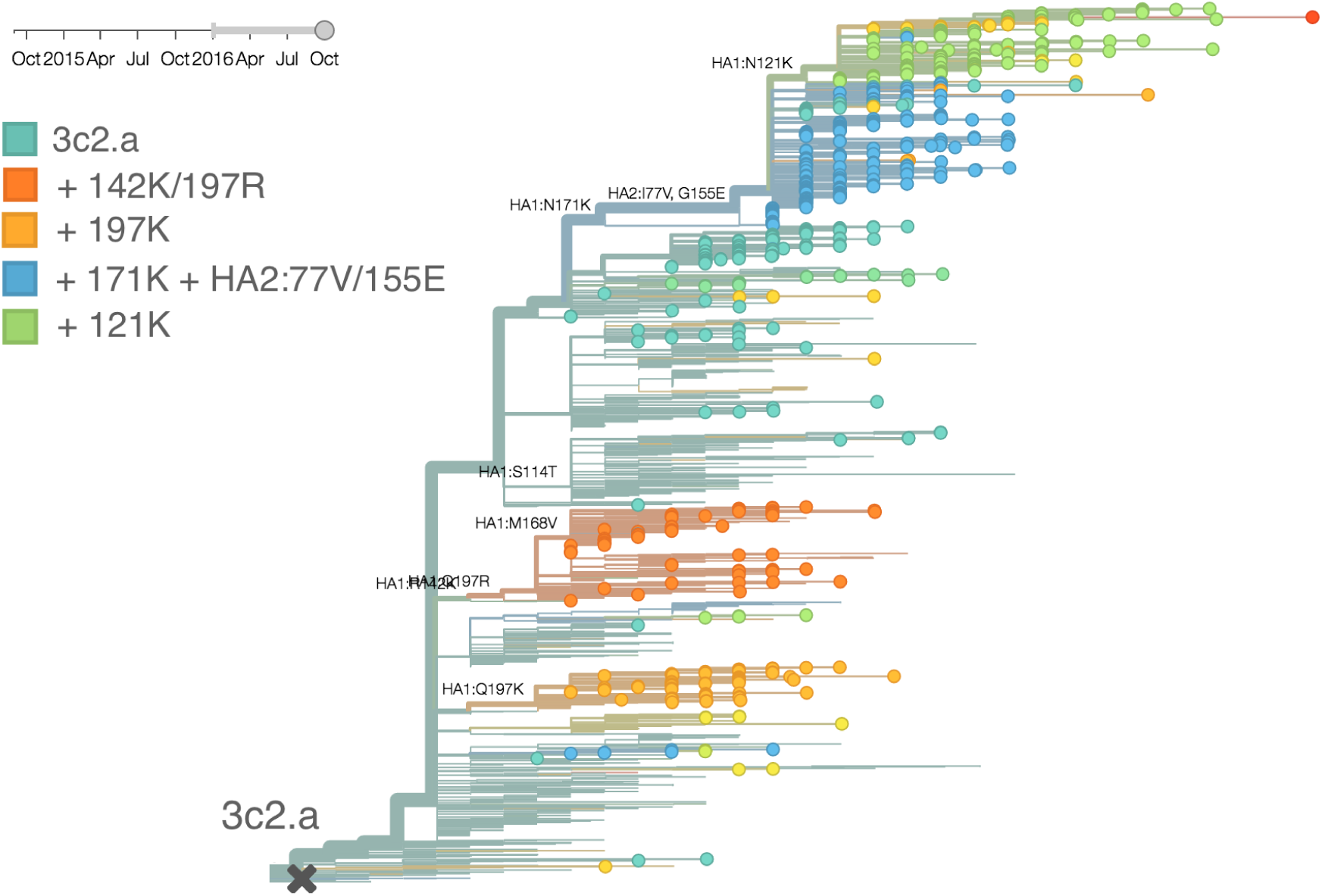
H3N2 / 3c2. a phylogeny colored by genotype.

The 142K/197R clade first emerged around June 2015 and rose to nearly 20% global frequency in Nov 2015 (Fig. 6). However, it’s declined throughout 2016. We doubt this is a competitive virus based on current clade success, but site 142 mutated several times within 3c2a to 142G or 142K suggesting that mutating this position might enable the virus to escape preexisting immunity at a fitness cost; it warrants continued observation. The 197K clade emerged in late 2015 and has slowly grown in frequency since. We estimate that it now comprises *∼*14% of H3N2 viruses. However, 197K is at extremely low frequency in Asia with an estimated present-day frequency *∼*2%. On the other hand, clade 171K viruses have done remarkably well throughout 2016 and now comprise an estimated 69% of currently circulating H3N2 viruses. This steady and rapid increase is strongly suggestive of an adaptive origin. Notably, 171K emerged and spread first in Asia, reaching nearly 80% frequency in April 2016. Higher frequency of 171K in Asia is expected to spread to the rest of the world given historic geographic observations [3]. Within 171K, the variant 121K has spread. However, it doesn’t appear to be spreading more rapidly than its parent 171K clade. Phylogenetic patterns suggest 171K as driver rather than 121K. Still, the rate of increase of 121K suggests that a sizable fraction of 2016-2017 viruses will be comprised of 121K.

*Without strong competition from another novel H3N2 virus, we believe that 171K will continue to increase in frequency in the global population and predominate in the 2016-2017 influenza season.* The continued spread of 171K is fully in line with the predictions we made in Feb 2016 [4]. At this point, we can’t say whether the 171K mutation in HA1 or the HA2 mutations 77V/155E (or some combination) is driving the selective spread of this clade.

**Figure 6.**
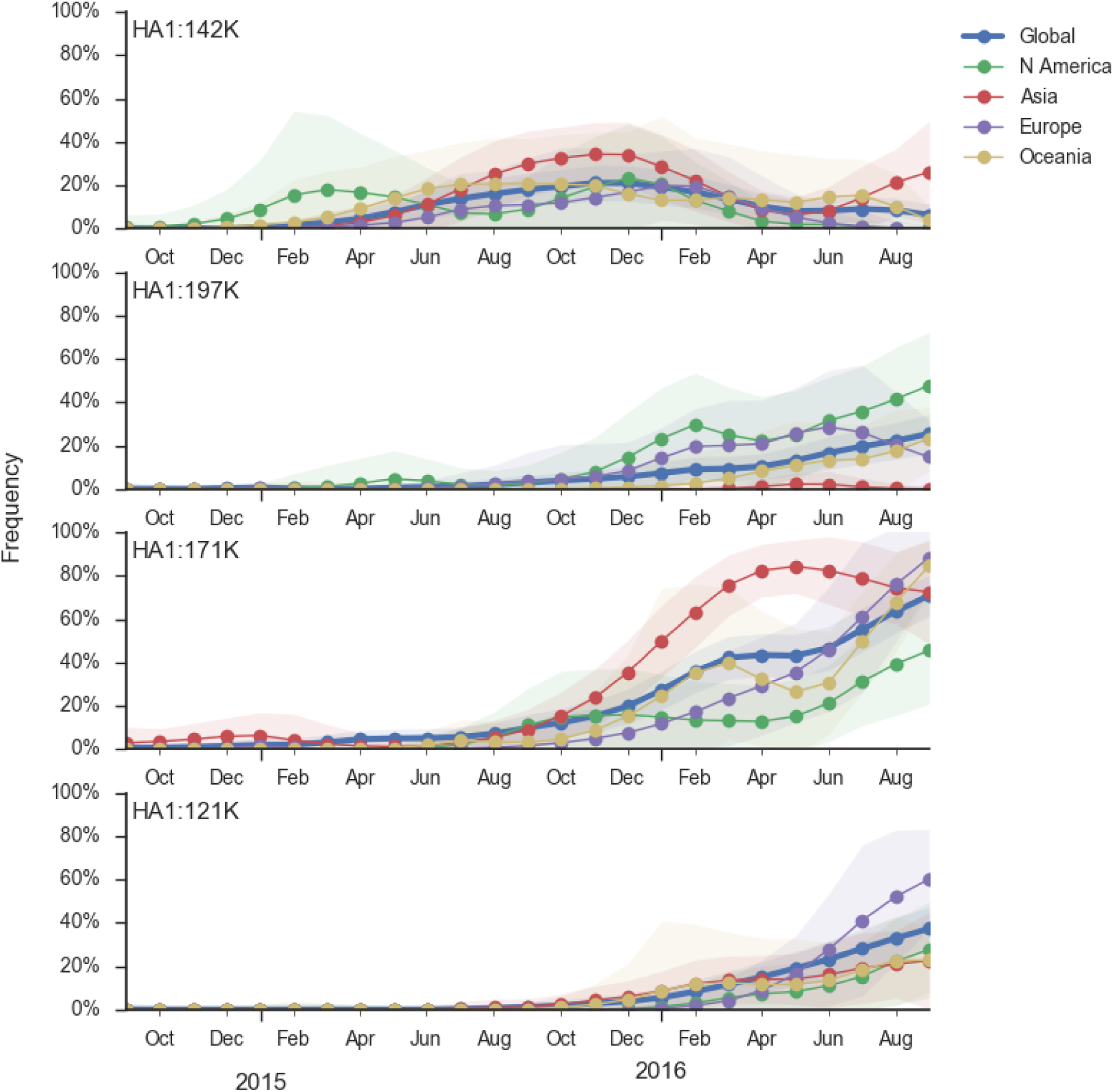
Frequency trajectories of 3c2.a subclades. We estimate frequencies of different clades based on sample counts and collection dates. We use a Brownian motion process prior to smooth frequencies from month-to-month. Transparent bands show an estimate the 95% confidence interval based on sample counts. The final point represents our frequency estimate for Sep 1 2016.

Other indicators suggest evolutionary success of 171K viruses. Notably “local branching index” [5] supports 171K as a high fitness virus (Fig. 7). 3c3.a viruses and other clades within 3c2.a and do not show signal in the “local branching index” analysis.

**Figure 7.**
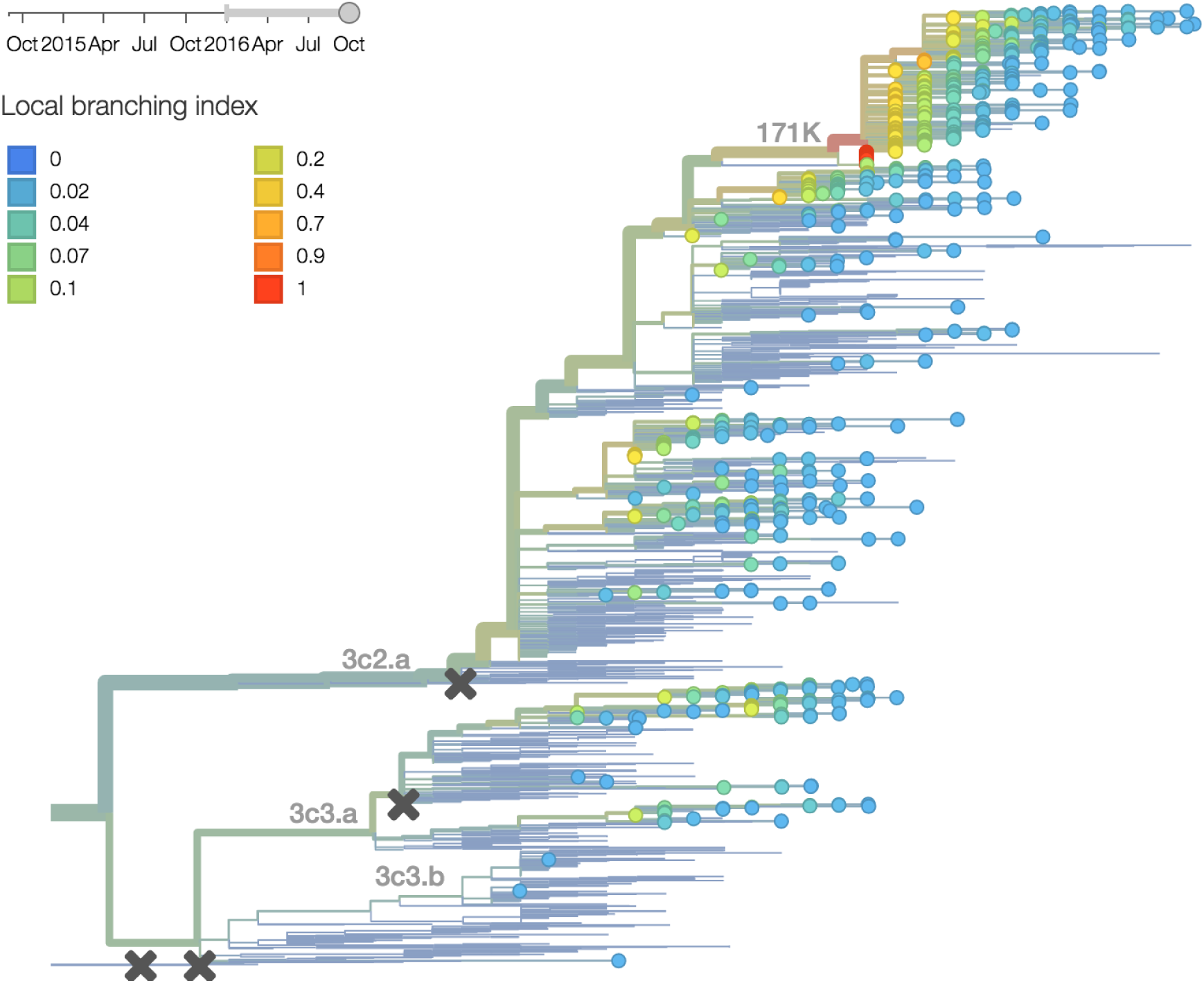
H3N2 phylogeny colored by local branching index.

Unfortunately, we lack sufficient recent serological data to distinguish antigenic evolution for most subclades within 3c2.a and 3c3.a. These observations derive entirely from genetic data.

There are a variety of viruses at the base of the 171K clade, possessing HA1:171K along with HA2:77V/155E, but lacking further amino acid changes in HA (Fig. 8).

**Figure 8.**
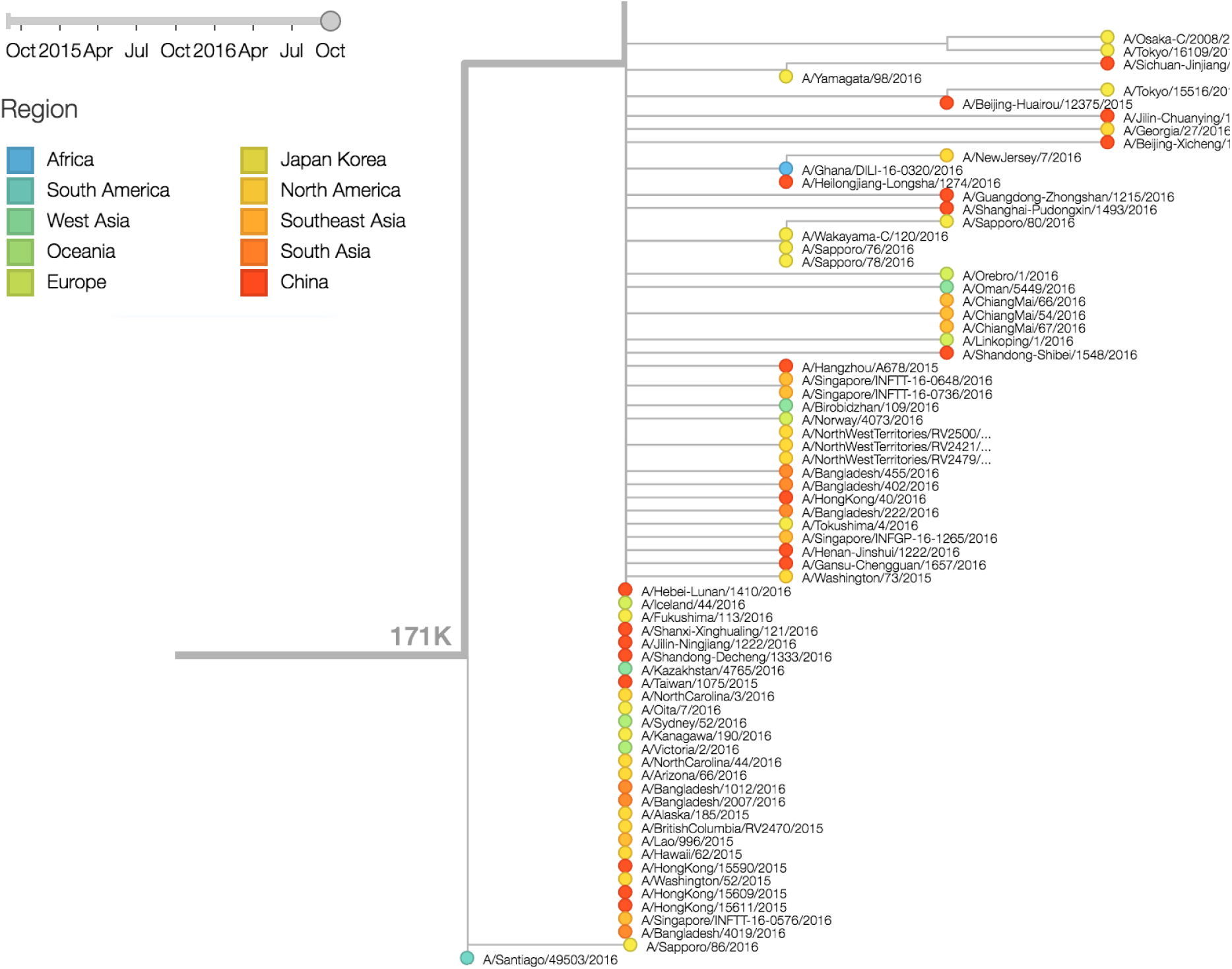
Viruses basal to the (HA1:171K, HA2:77V/155E) clade. Viruses are colored according to sampling region.

## A/H1N1pdm

**The clade 6b.1, comprising 84N/162N/216T, has continued to rise and predominate in the H1N1pdm population throughout 2016. Almost all circulating H1N1pdm viruses are now 6b.1. There is not yet obvious evolution within this clade.**

As discussed above, we base our primary analysis on a set of viruses collected between Sep 2014 and Aug 2016, comprising approximately 100 viruses per month where available and seeking to equilibrate sample counts geographically where possible (Fig. 9). Recent months through June 2016 have largely sufficient sample counts and sample distributions. There are fewer samples from July to present.

**Figure 9.**
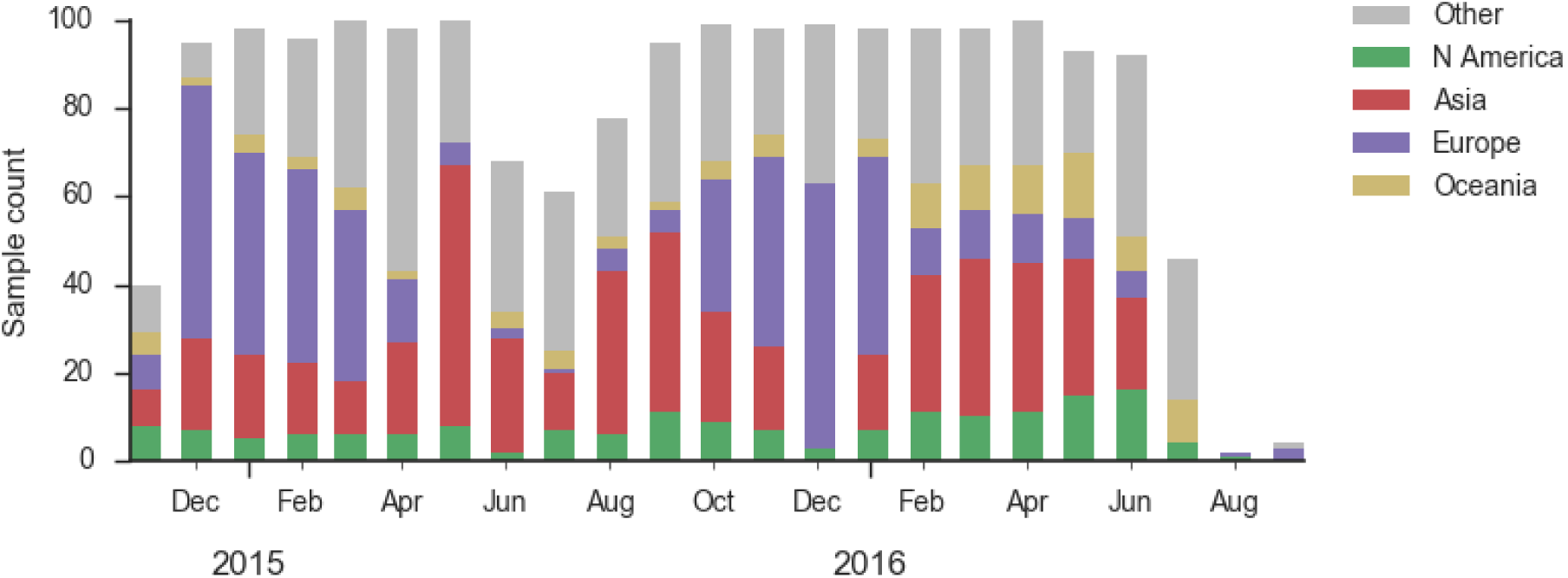
Sample counts through time and across regions. This is a stacked bar plot, so that in good months there are *∼*100 total samples and *∼*15 samples each from North America and from Europe.

Within clade 6b, two major genetic variants have emerged (Fig. 10). These are clade 6b.1 comprised of HA1:84N/162N/216T and clade 6b.2 comprised of HA1:152T and HA2:174E. Most recent samples have been from 6b.1 viruses, with 85% of 2016 samples being from 6b.1 viruses.

**Figure 10.**
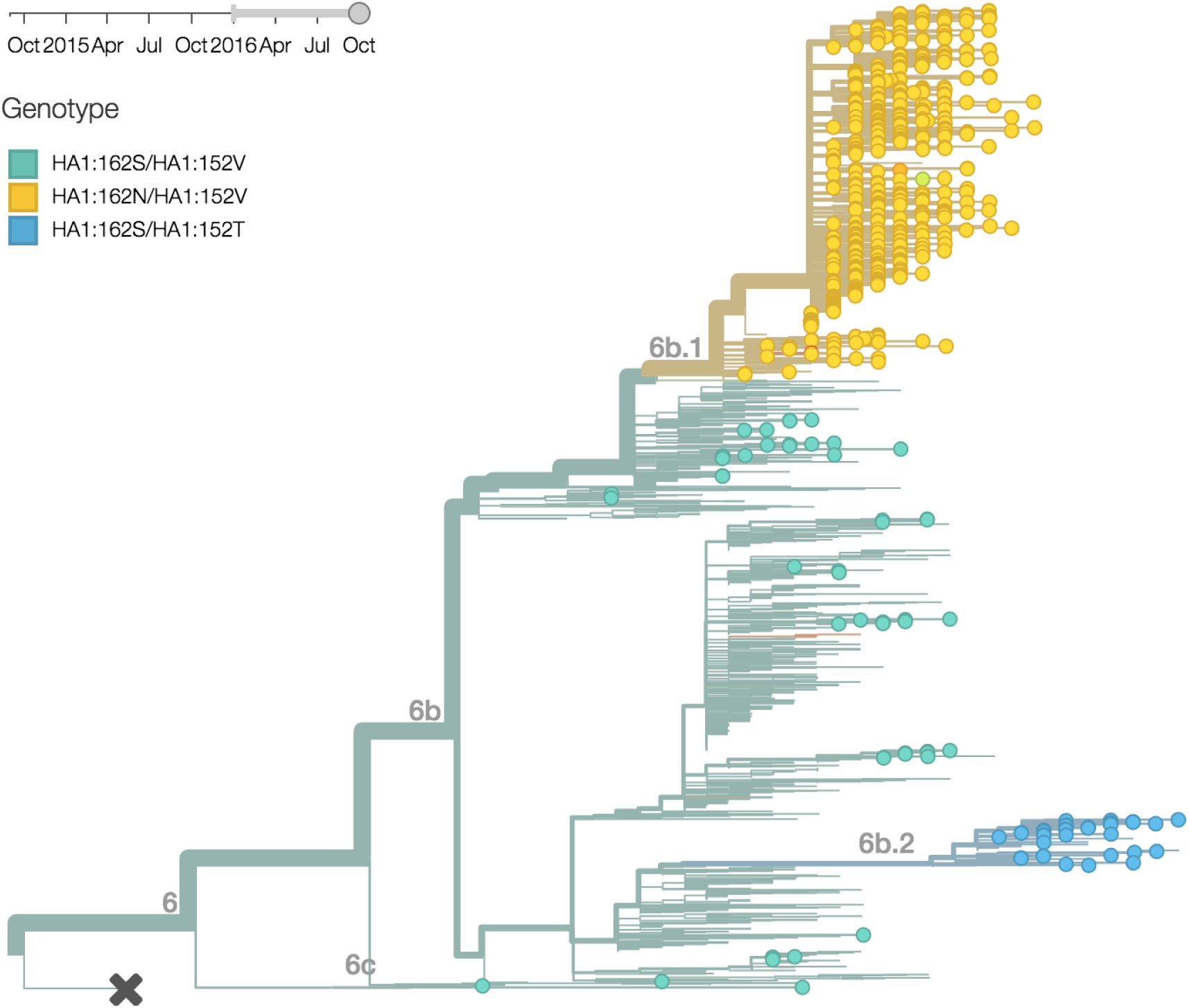
H1N1pdm phylogeny colored by genotype.

At this point, all regions of the world are dominated by 6b.1 viruses (Fig. 11). This clade rose from low frequency in Aug 2015 to reach present day global frequencies of *∼*98%. There remain a minority of circulating 6b.2 viruses. We estimate 2% of H1N1pdm viruses globally to be 6b.2. These are slightly higher prevalence in Asia, but still a distinct minority. We estimate that 6b.1 is at 85% frequency in Asia, while 6b.2 is at 15% in Asia. Notably, the frequency of 6b.2 has remained stable for almost 12 months.

*Every indication suggests the continued dominance of 6b.1 viruses, and their extremely rapid rise suggests a selective origin. We are now watching for the emergence of genetic variants within the 6b.1 clade.* Notably, the continued rise and dominance of 6b.1 viruses fits with our predictions from Feb 2016 [4].

**Figure 11.**
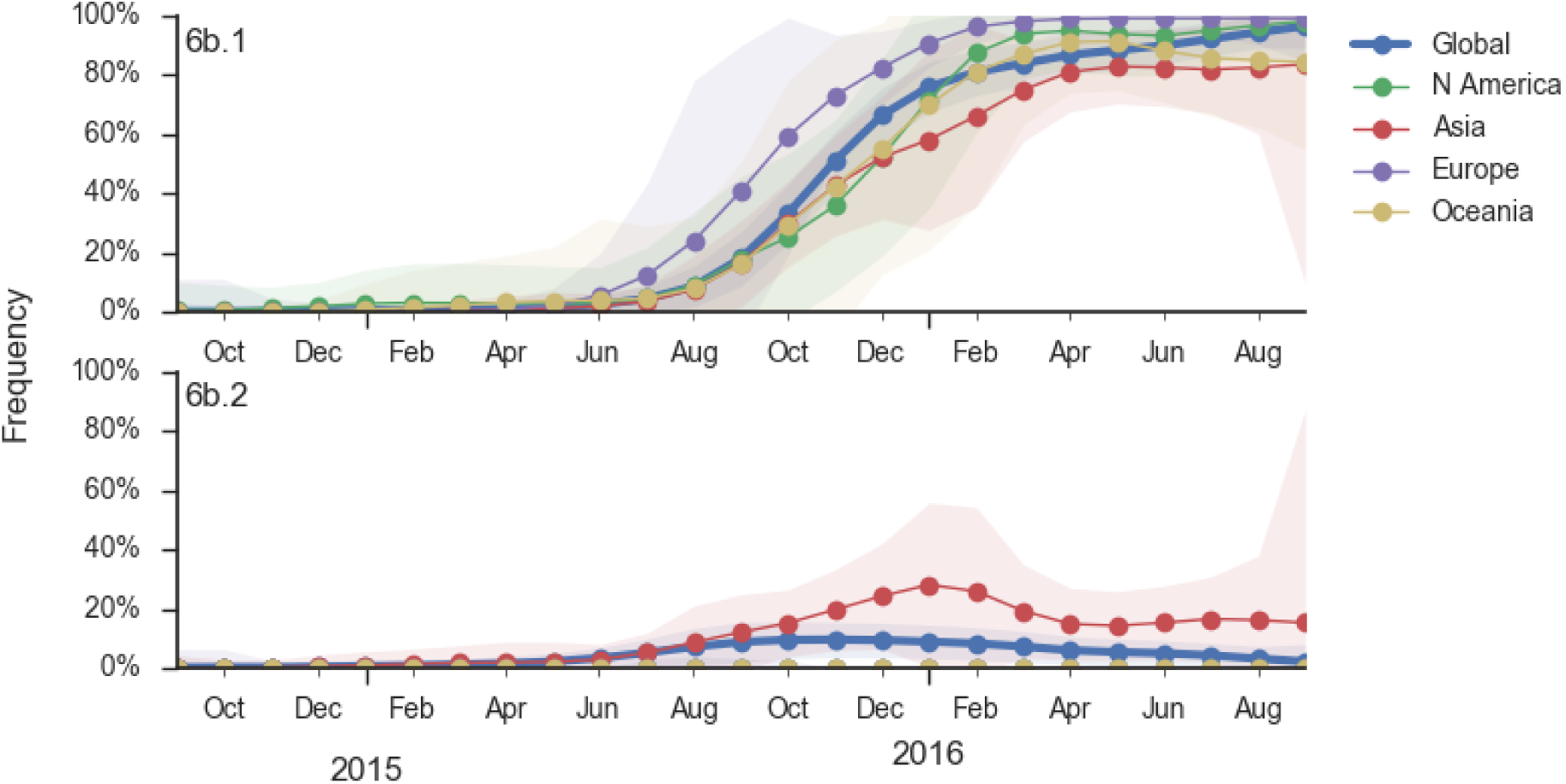
Frequency trajectories of H1N1pdm clades. We estimate frequencies of different clades based on sample counts and collection dates. We use a Brownian motion process prior to smooth frequencies from month-to-month. Transparent bands show an estimate the 95% confidence interval based on sample counts. The final point represents our frequency estimate for Sep 1 2016.

Analysis of local branching index [5] also flags the clade 6b.1 as rapidly expanding and the most successful lineages within H1N1pdm (Fig. 12).

**Figure 12.**
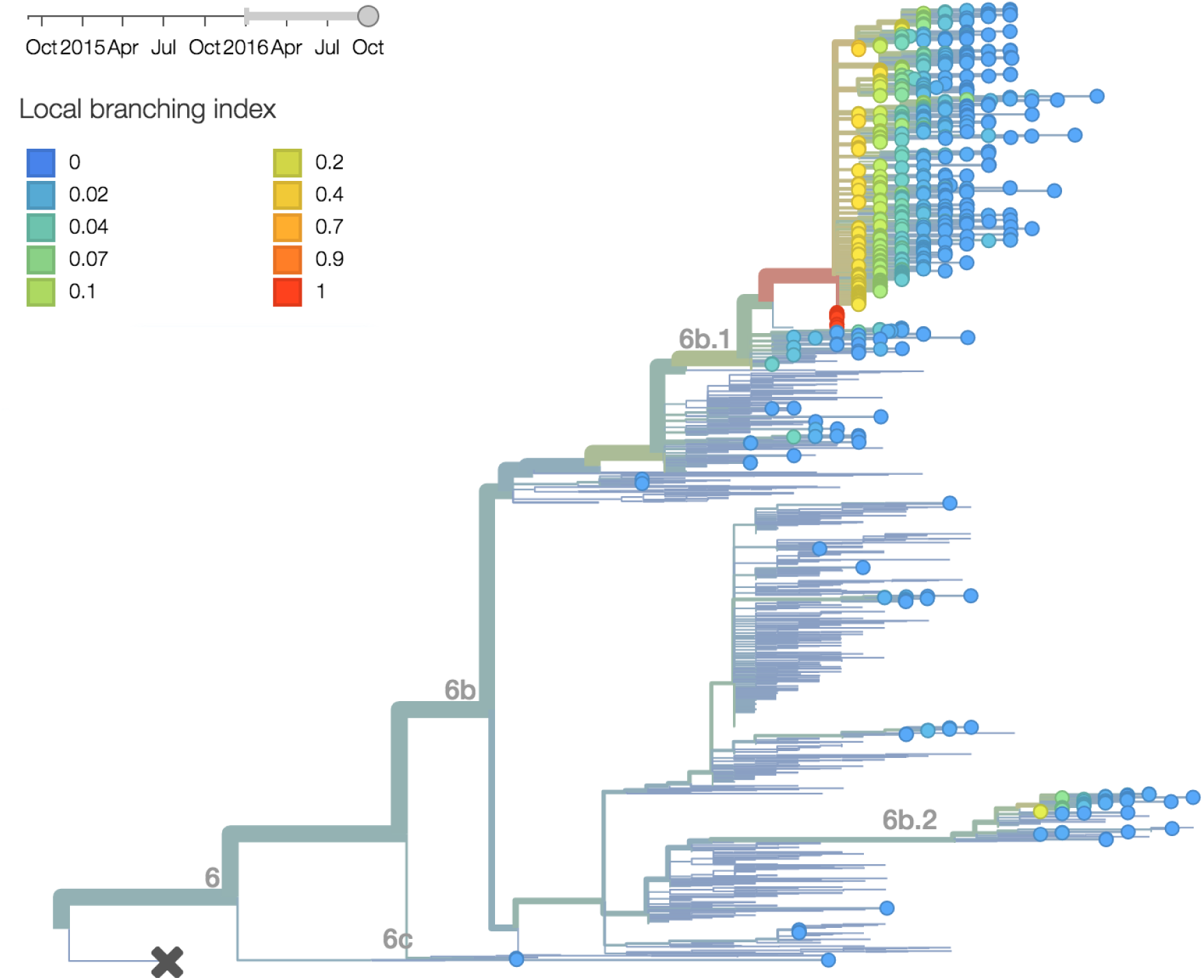
H1N1pdm phylogeny colored by local branching index.

Analysis of antigenic evolution via the method of Neher et al. (2015) [6] and using recent HI data from the Feb 2016 VCM report [7] from the Crick Institute Collaborating Center suggests a small effect on HI titer in both the 6b.1 and 6b.2 clades (Fig. 13). In this case, 6b.1 shows an antigenic impact of 0.65 log2 units and 6b.2 shows an impact of 0.66 log2 units. Interesting, this analysis places the impact on clade 6b.1 from the 84N mutation rather than the later 162N or 216T mutations.

**Figure 13.**
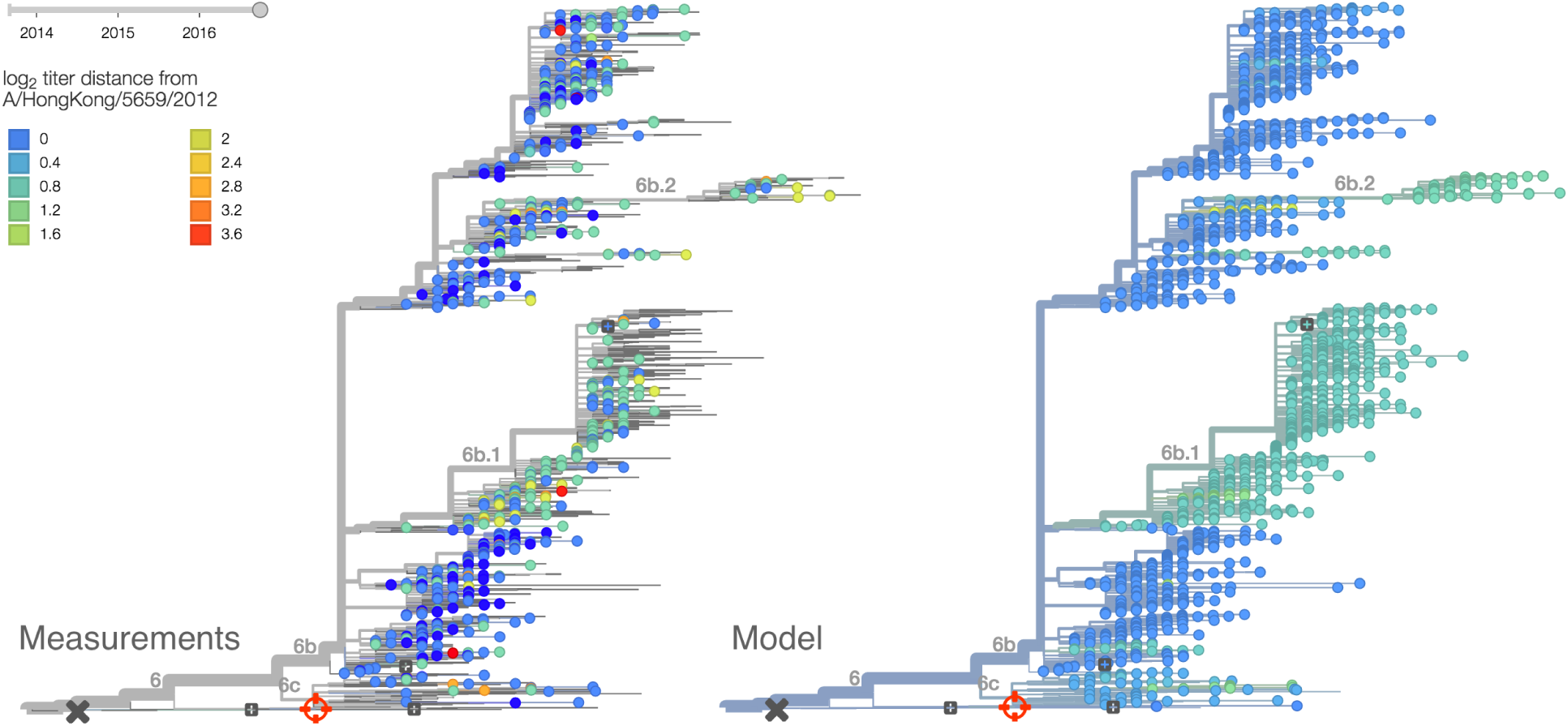
H1N1pdm phylogeny colored by antigenicity. The left panel shows individual HI measurements of viruses against serum from A/HongKong/5659/2012. Cooler color indicates great antigenic similarity (less titer drop going from homologous to heterologous titers). The right panel shows model estimates of antigenic divergence from A/HongKong/5659/2012. These estimates combine HI measurements from many different sera. Data is current through Feb 2016.

## B/Vic

**Within clade 1A viruses, the clade 129D/146I/117V has risen to high frequency, but at a rate that suggests a smaller effect of natural selection. At this point, 117V viruses are dominant in the global viral population.**

As above, we base our primary analysis on a set of viruses collected between Sep 2014 and Aug 2016, comprising approximately 100 viruses per month where available and seeking to equilibrate sample counts geographically where possible (Fig. 14).

**Figure 14.**
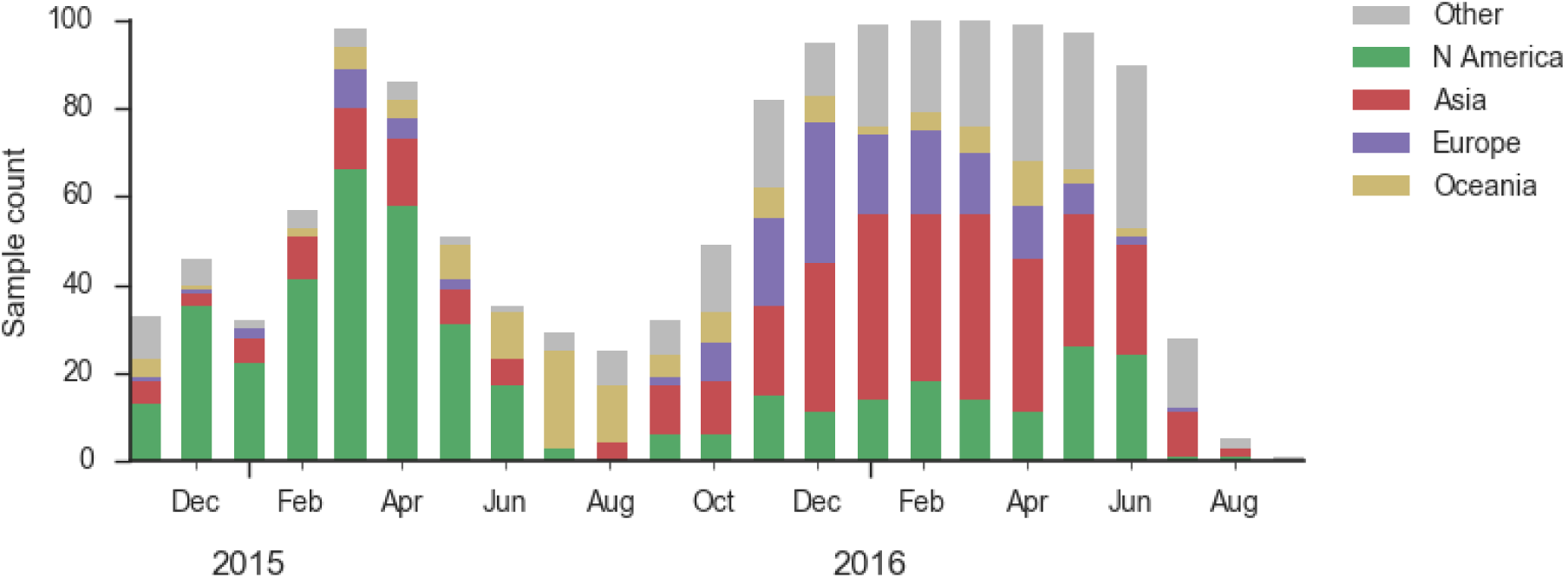
Sample counts through time and across regions. This is a stacked bar plot, so that in good months there are *∼*100 total samples and *∼*15 samples each from North America and from Europe.

In the past year, B/Vic clade 129D/146I/117V has dominated the viral population, with 94% of samples possessing the 117V mutation (Fig. 15). This dominance has increased throughout 2016 and we estimate that currently circulating Vic viruses are 94% 117V. At this point, we haven’t observed new variants of appreciable frequency within the 117V clade and there aren’t decent competitors outside the 117V clade.

**Figure 15.**
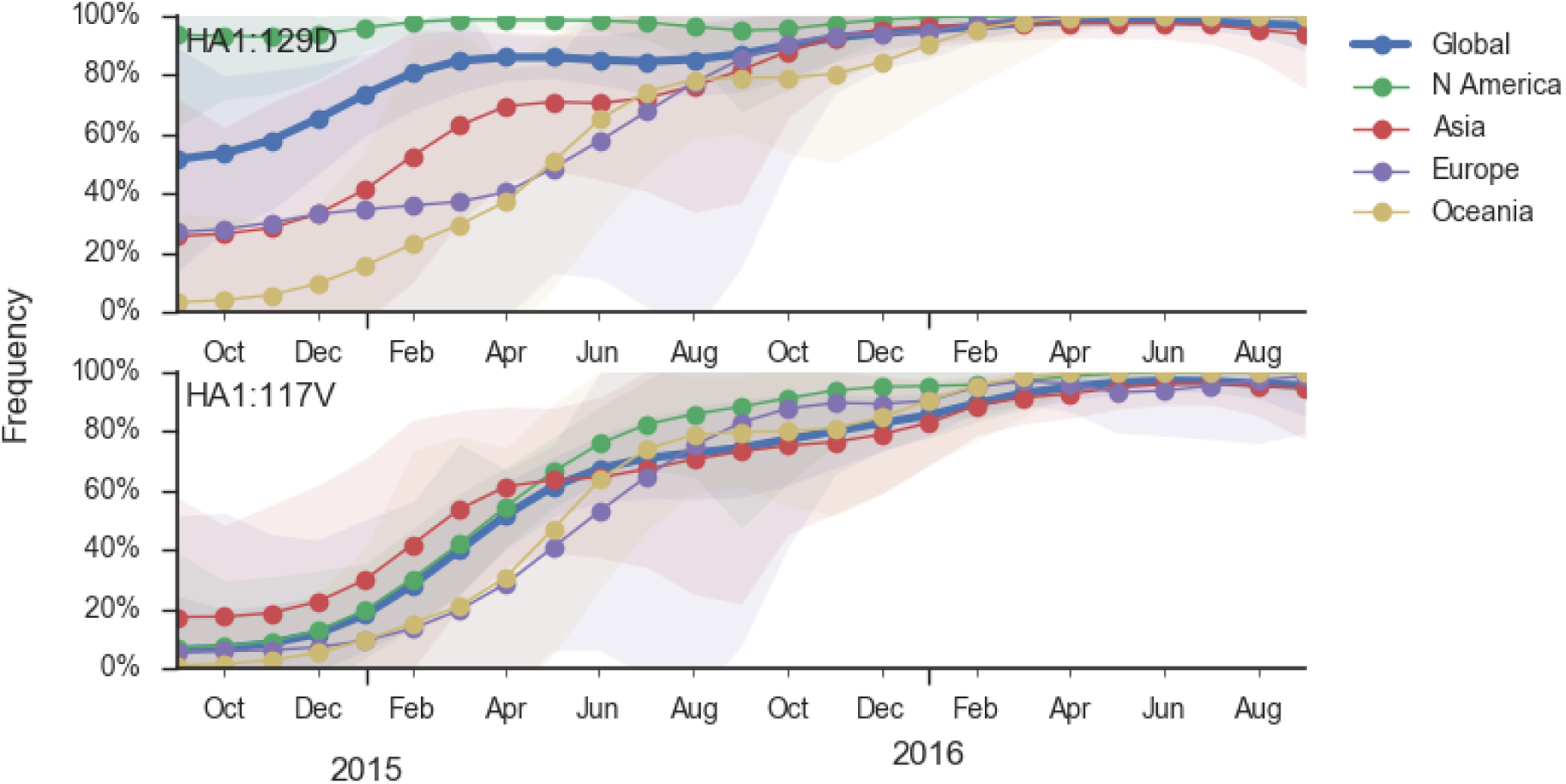
Frequency trajectories of B/Vic variants. We estimate frequencies of different clades based on sample counts and collection dates. We use a Brownian motion process prior to smooth frequencies from month-to-month. Transparent bands show an estimate the 95% confidence interval based on sample counts. The final point represents our frequency estimate for Sep 1 2016.

Additionally, the 117V clade has been growing most quickly and is picked by the local branching index [5] as the currently most successful clade (Fig. 16). *All indicators suggest the continued success of 117V in the coming year.*

**Figure 16.**
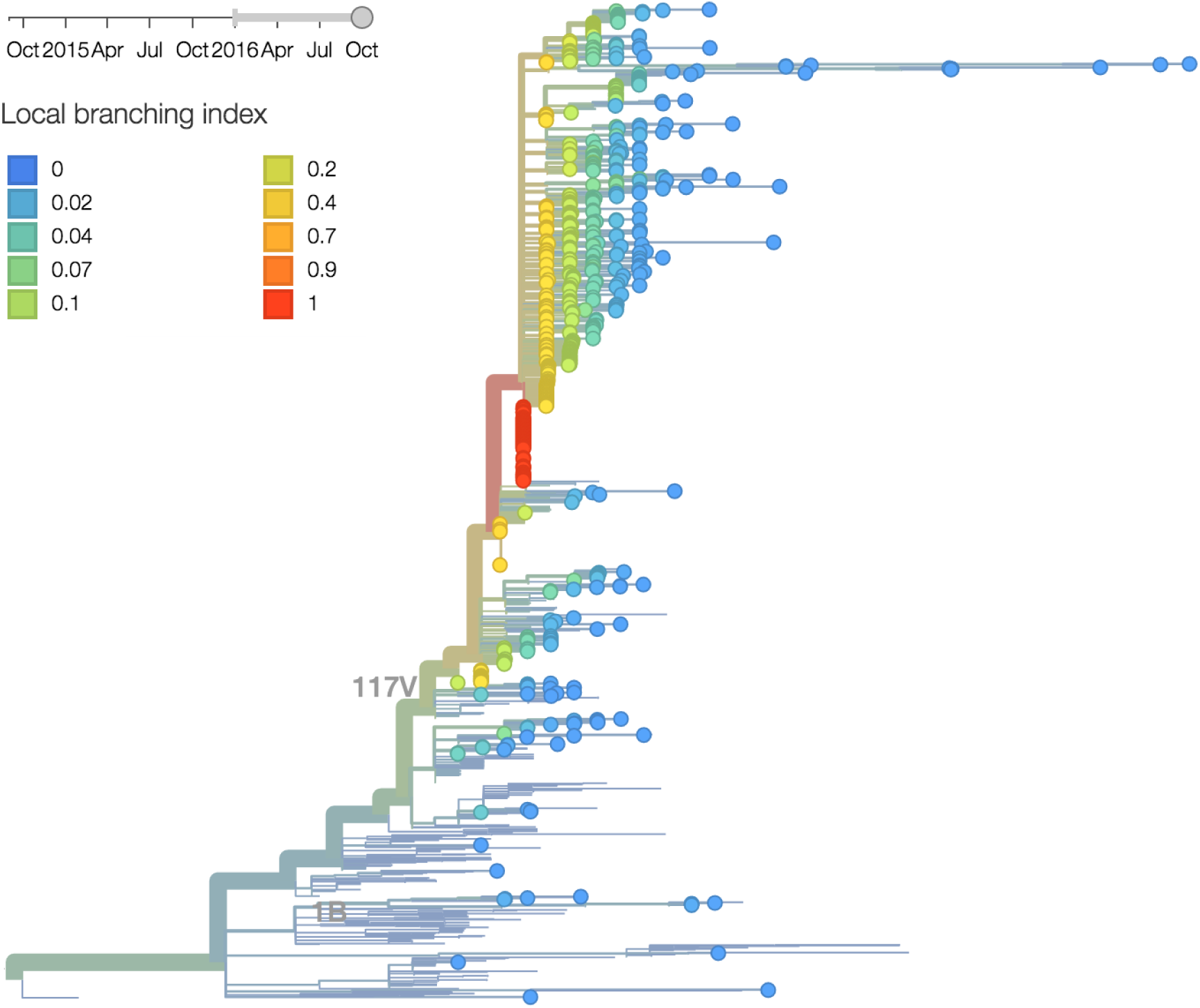
B/Vic phylogeny colored by local branching index.

## B/Yam

**Clade 3 has continued to predominate the B/Yamagata population. Within clade 3 the 172Q/251V subclade has risen to predominate. There is limited diversity within 172Q/251V viruses, with the largest subclade comprised of 211R viruses.**

As above, we base our primary analysis on a set of viruses collected between Sep 2014 and Jul 2016, comprising approximately 100 viruses per month where available and seeking to equilibrate sample counts geographically where possible (Fig. 17).

**Figure 17.**
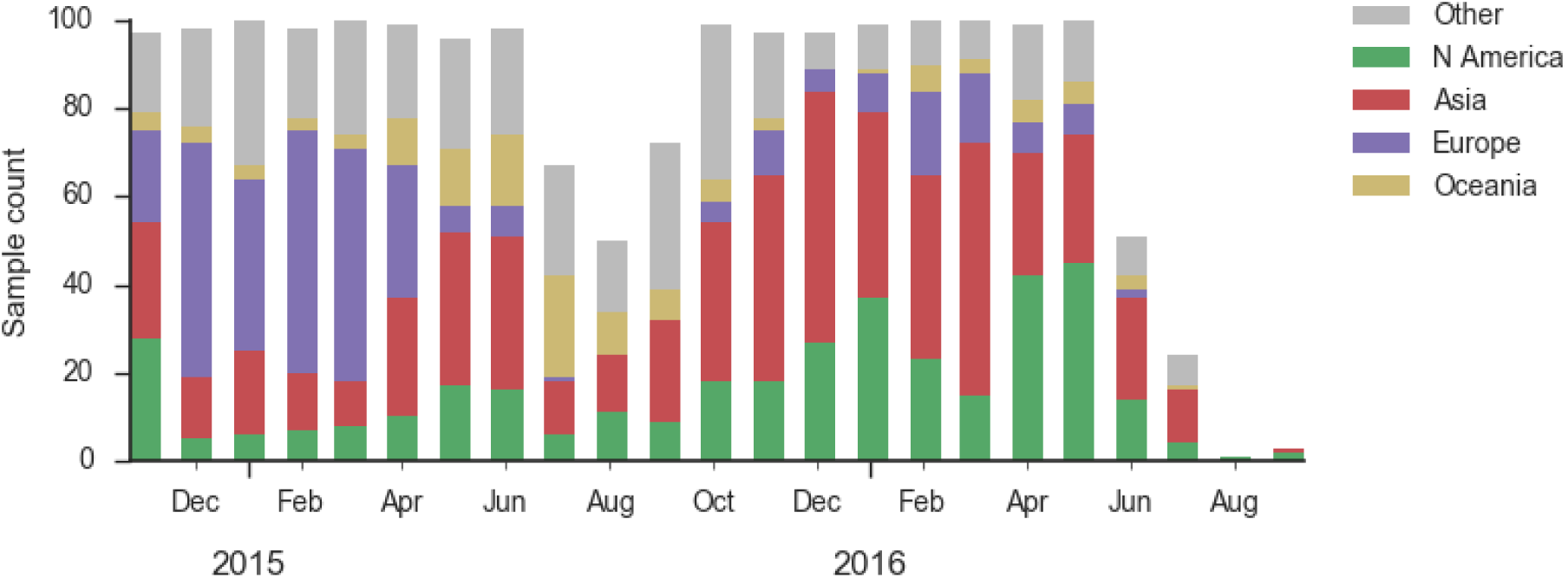
Sample counts through time and across regions. This is a stacked bar plot, so that in good months there are *∼*100 total samples and *∼*15 samples each from North America and from Europe.

During 2016, the vast majority of B/Yamagata isolates were of clade 3 viruses (Fig. 18). We estimate that the current frequency of clade 2 is now *∼*2%. At this rate, we expect clade 2 to go extinct in the coming year.

**Figure 18.**
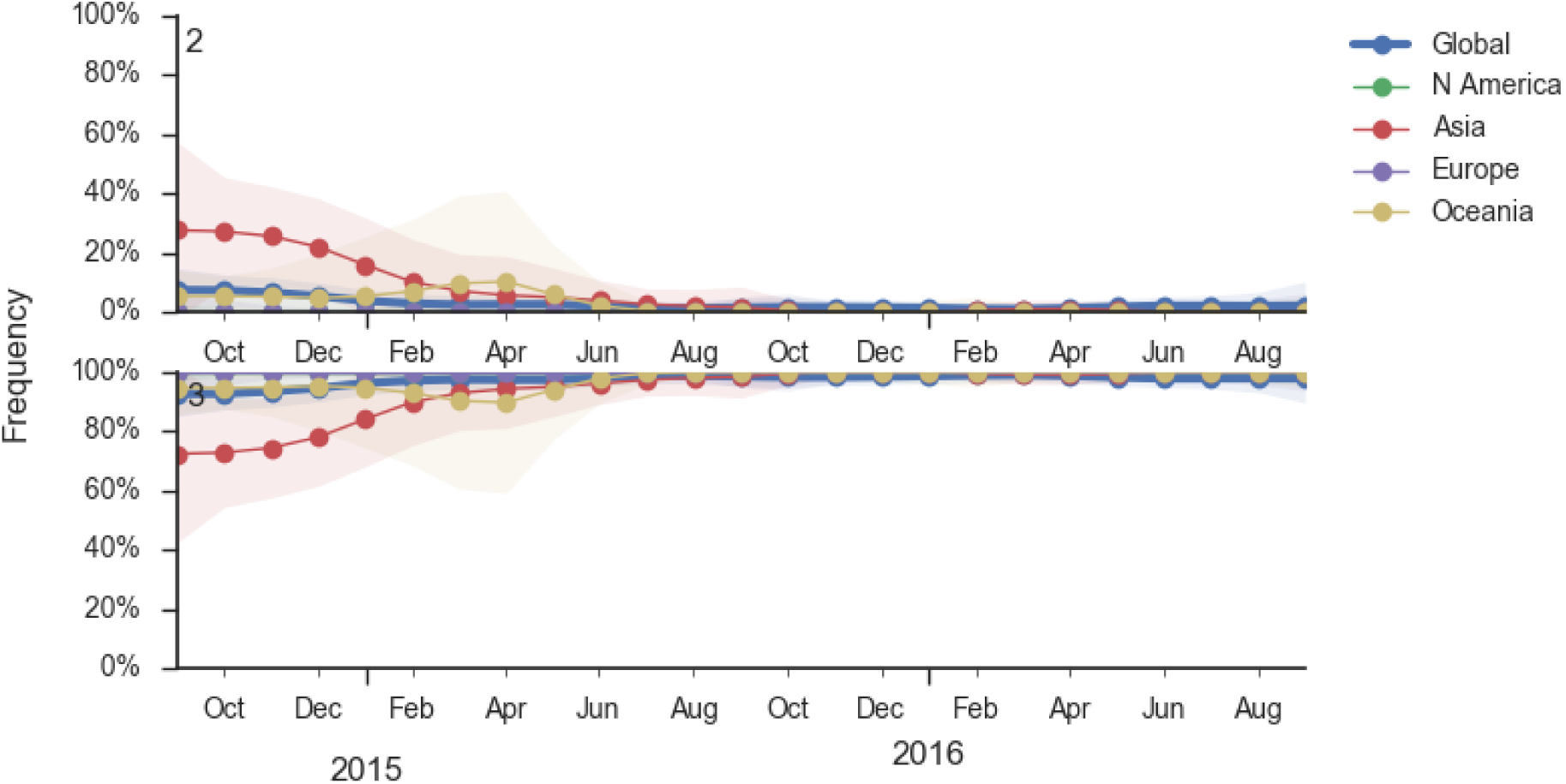
Frequency trajectories of B/Yam clades. We estimate frequencies of different clades based on sample counts and collection dates. We use a Brownian motion process prior to smooth frequencies from month-to-month. Transparent bands show an estimate the 95% confidence interval based on sample counts. The final point represents our frequency estimate for Sep 1 2016.

Within clade 3, HA1:172Q has predominated throughout 2015 and 2016 (Fig. 19). On this background the HA1:251V variant has emerged and on top of 251V, the HA1:211R variant has appeared.

**Figure 19.**
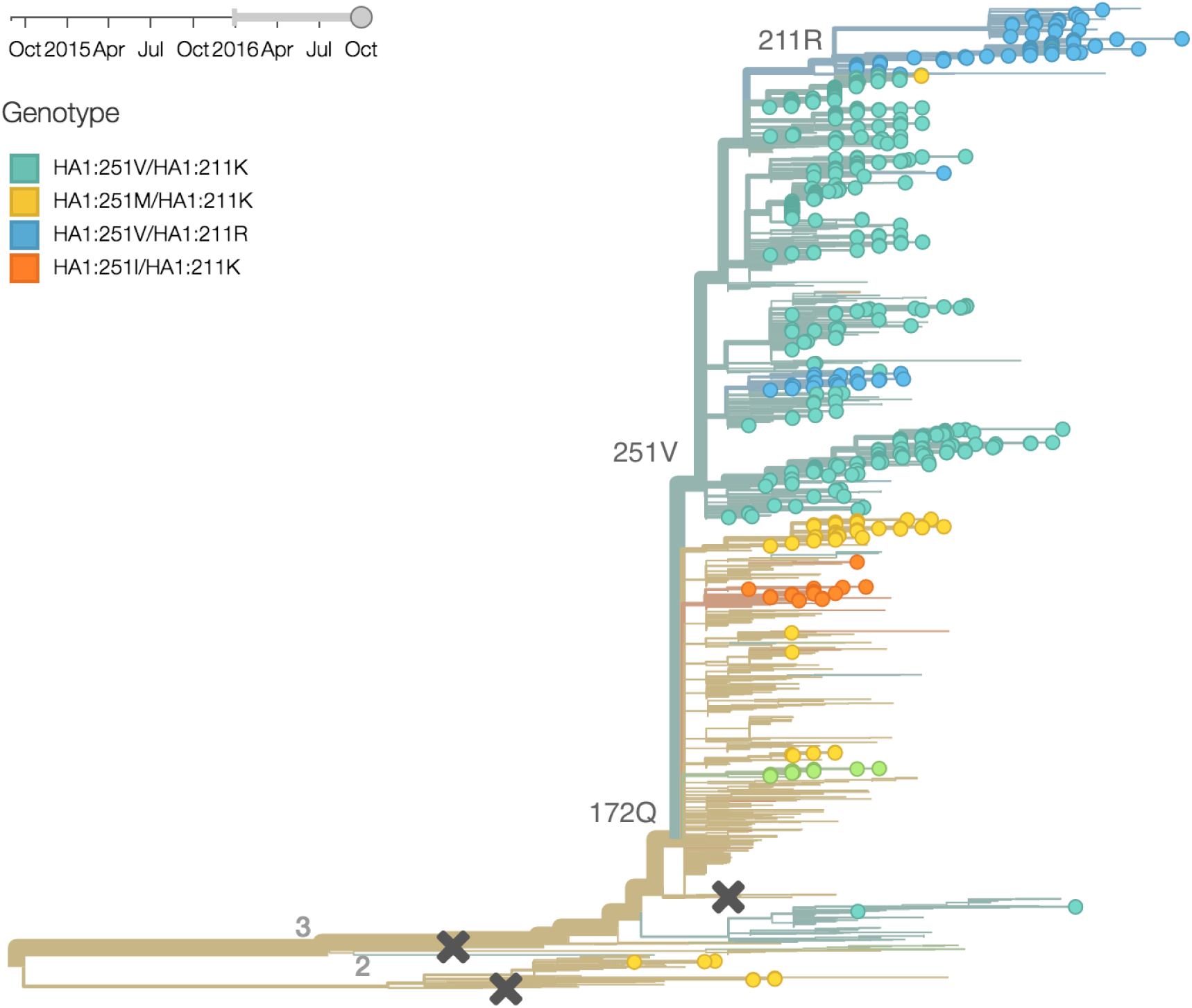
B/Yam phylogeny colored by genotype.

The 251V clade increased from low frequency in Oct 2014 to predominate in the population (Fig. 20). We estimate that 251V is currently at 79% globally. The 211R variant rose from low frequency in Apr 2015 to reach 34% in currently circulating viruses. The rate of increase, however, has been rather mild.

**Figure 20.**
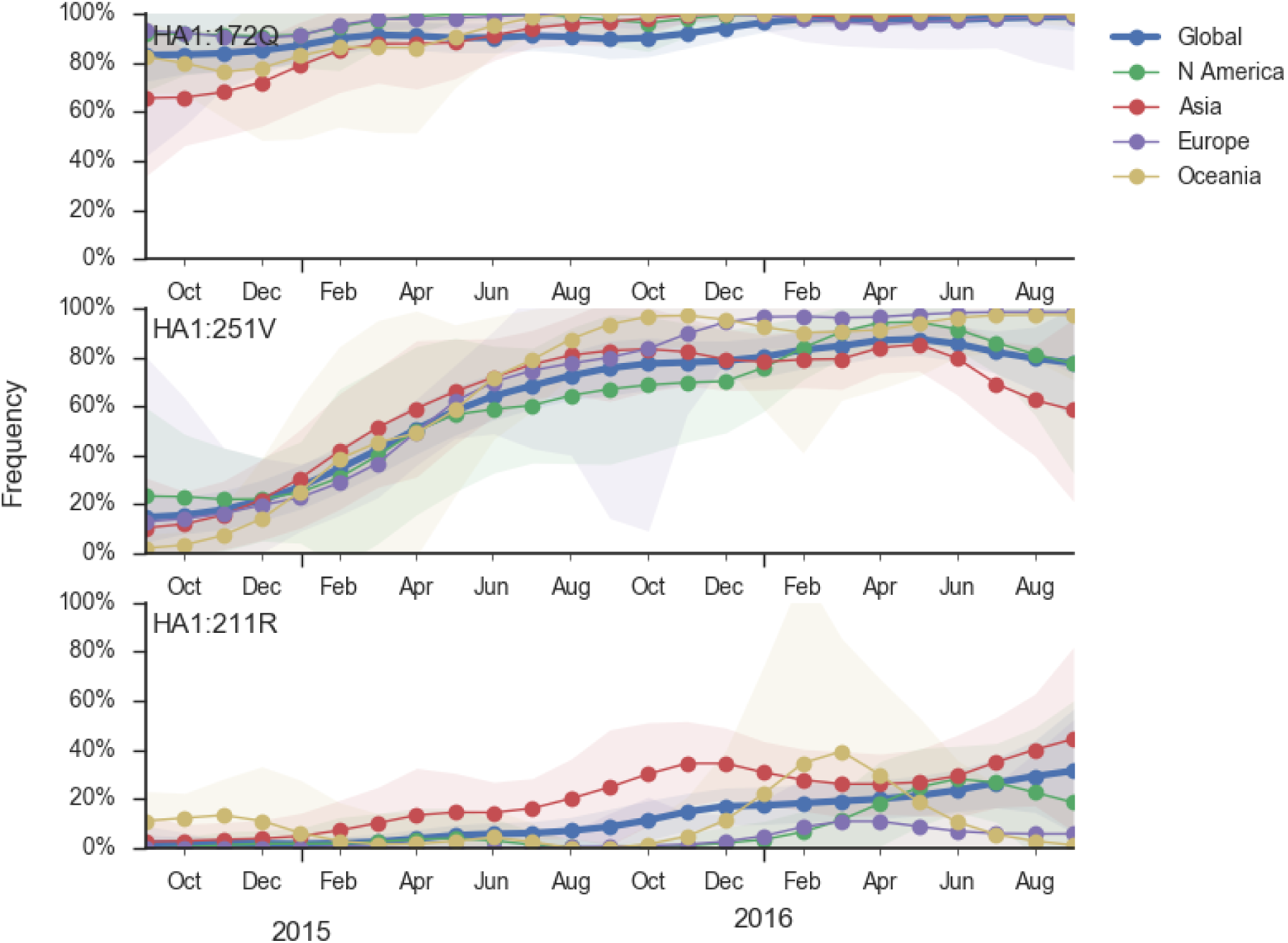
Frequency trajectories of B/Yam variants. We estimate frequencies of different clades based on sample counts and collection dates. We use a Brownian motion process prior to smooth frequencies from month-to-month. Transparent bands show an estimate the 95% confidence interval based on sample counts. The final point represents our frequency estimate for Sep 1 2016.

## A/H3N2 (all sequences)

The clade and mutation frequencies discussed above were based on a limited sequence subsample with an equitable geographical distribution. More accurate region specific trajectories can be estimated using all sequence data available in GISAID. We repeated the clade and mutation frequency estimation using up to 500 sequences per month and region (Fig. 21).

**Figure 21.**
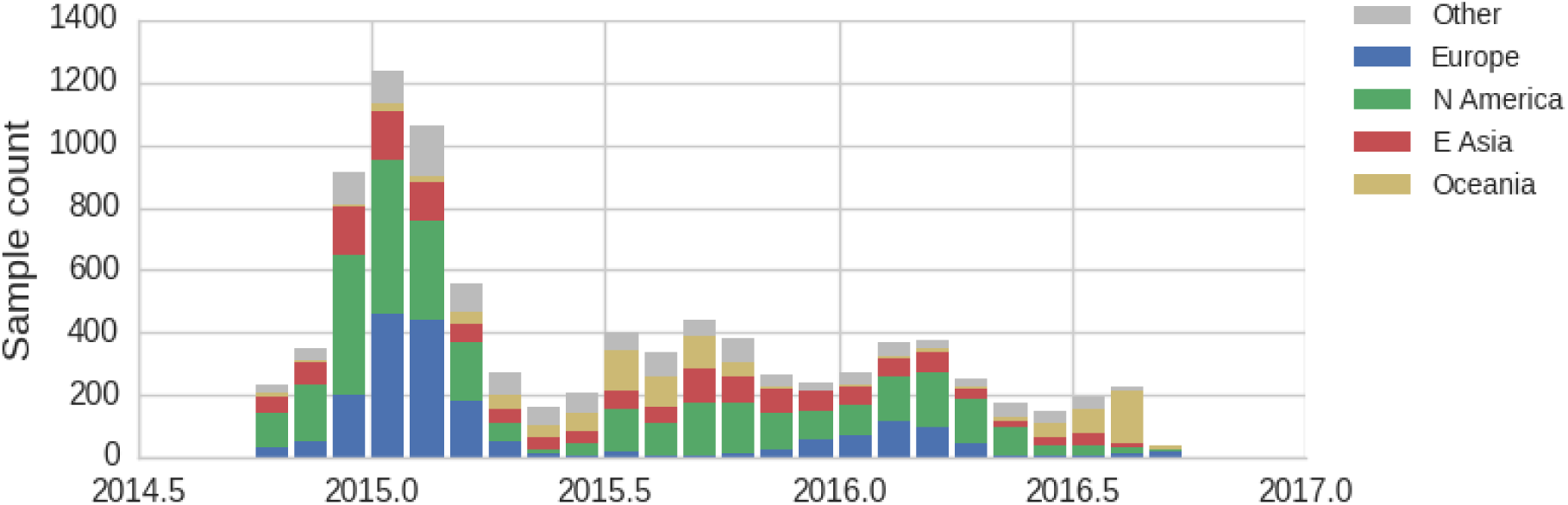
Sample counts through time and across regions. This is a stacked bar plot, so that in the peak of the 2014-2015 season there are *∼*700 total samples per month.

With this more inclusive sampling, northern hemisphere winter months tend to be dominated by North America and Europe, while other months are dominated by samples from North America, Asia, and Oceania (Fig. 22). The global average will track the regions contributing the majority of the sequence data.

**Figure 22.**
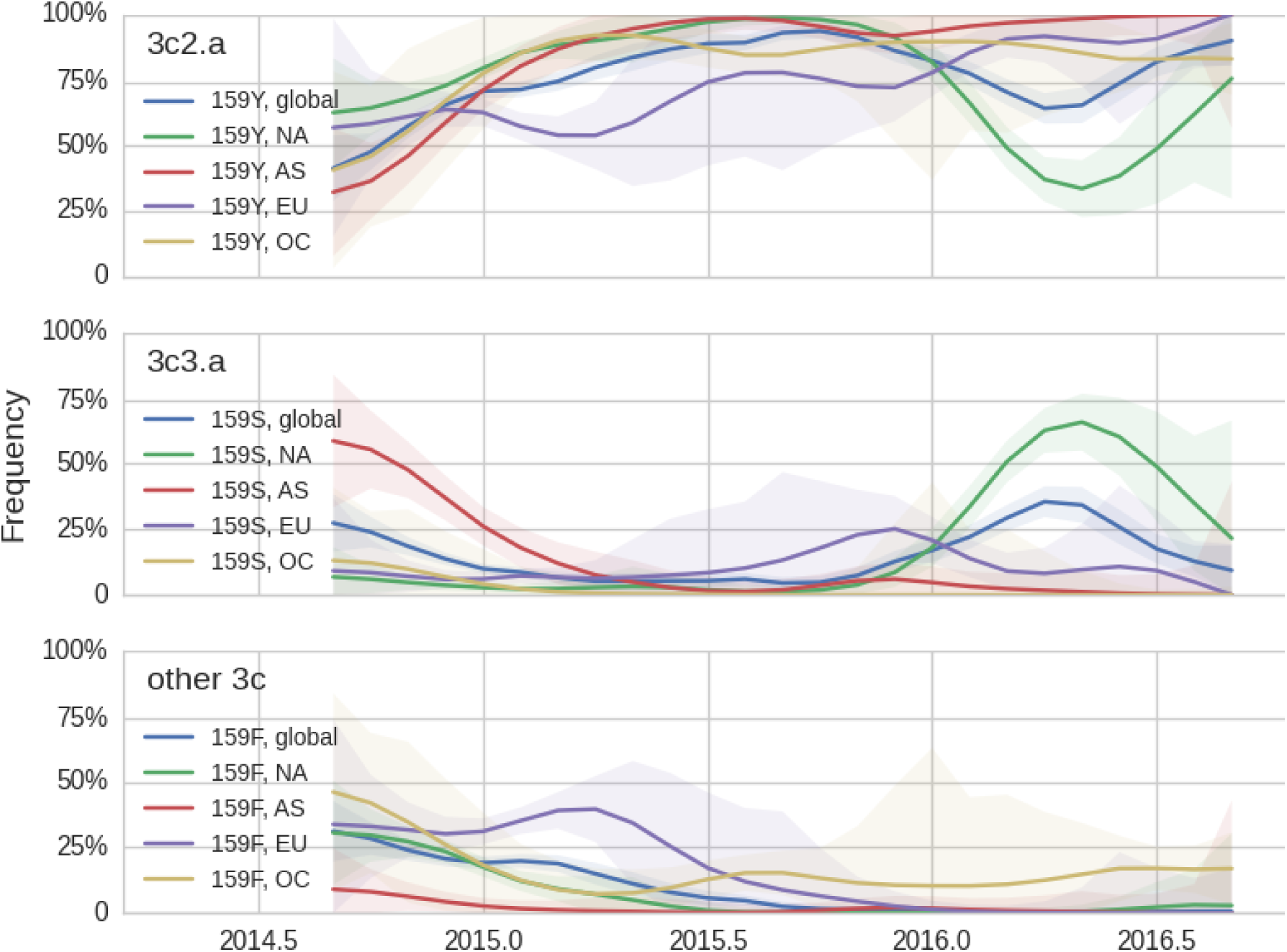
Frequency trajectories of H3N2 clades. We estimate frequencies of different clades based on sample counts and collection dates. We use a Brownian motion process prior to smooth frequencies from month-to-month. Transparent bands show an estimate the 95% confidence interval based on sample counts. The final point represents our frequency estimate for Sep 1 2016.

The H3N2 clades 3c3.a (159S) and 3c2.a (159Y) show broadly similar trajectories as discussed above (Fig. 23). 3c2.a came to dominate Asia completely, while viruses outside of 3c3.a and 3c2.a continue to circulate at low levels in Oceania. Within 3c2.a, the mutation 171K dominates across all regions.

**Figure 23.**
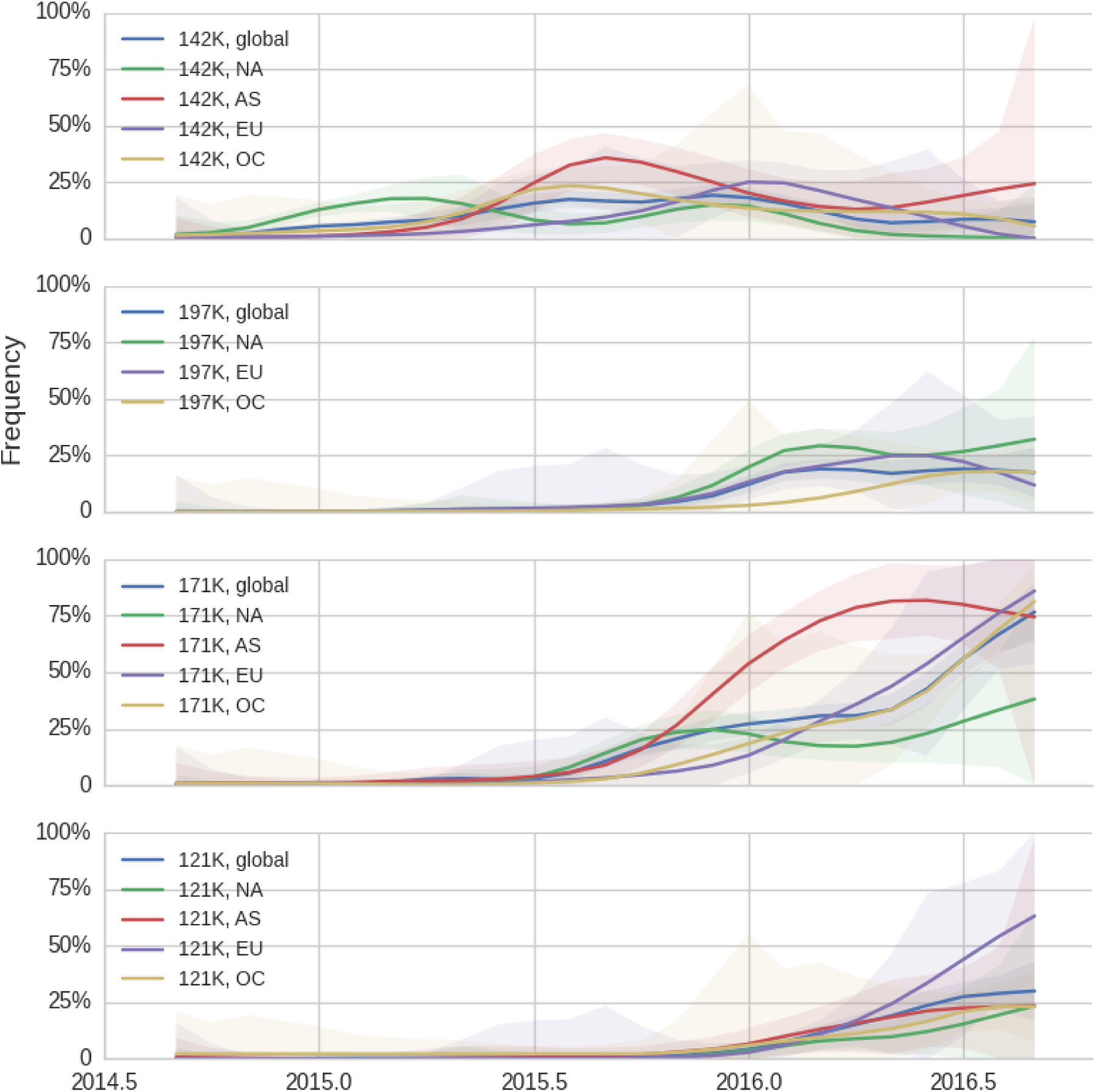
Frequency trajectories of H3N2 mutations. We estimate frequencies of different clades based on sample counts and collection dates. We use a Brownian motion process prior to smooth frequencies from month-to-month. Transparent bands show an estimate the 95% confidence interval based on sample counts. The final point represents our frequency estimate for Sep 1 2016.

